# Mitotic spindle disassembly in human cells relies on CRIPT-bearing hierarchical redox signals

**DOI:** 10.1101/869123

**Authors:** Kehan Xu, Lingling Yang, Xiu Cheng, Xiaoyan Liu, Hao Huang, Haibing Tang, Xuejiao Xu, Jingyu Wang, Anan Jiang, Chunxue Wang, Meifang Lu, Zhengmei Lv, Lin Shen, Feifei Song, Haoqian Zhou, Haisheng Zhou, Xinhua Liu, Weibing Shi, Jinghua Zhou, Xuejun Li, Hong Li, Chunlin Cai

**Author notes:** These authors contributed equally to this work (Lingling Yang, Xiu Cheng, Xiaoyan Liu, Hao Huang).

## Abstract

Swift and complete spindle disassembly is essential for cell survival, yet how it happens is largely unknown. Here we used real-time live-cell microscopy and biochemical assays to show that a cysteine-rich protein CRIPT dictates the spindle disassembly in a redox-dependent manner in human cells. This previously reported cytoplasmic protein was found to have a confined nuclear localization during interphase but was distributed to spindles and underwent redox modifications to form disulfides within CXXC pairs during mitosis. Then, it interacts with and transfers redox response to tubulin subunits to induce microtubule depolymerization. The mutants with any of cysteine substitution completely block the spindle disassembly generating two cell populations with long-lasting metaphase spindles or spindle remnants. The live cell recordings of a disease-relevant mutant (CRIPT^C3Y^) revealed that microtubule depolymerization at spindle ends during anaphase and the entire spindle dissolution during telophase may share a common CRIPT-bearing redox-controlled mechanism.

## Introduction

The mitotic spindle is a highly dynamic structure undergoing continuous turnover between the assembly and disassembly. The assembly has been well studied, but how spindles disassemble is poorly understood especially in mammalian cells. Spindle disassembly occurs in a phase-specific way (Asbury, 2017). During anaphase, spindles undergo partial microtubule (MT) disassembling at their ends as a main mechanism for the poleward movement to separate the chromosomes. During telophase, spindles must be rapidly and entirely disassembled after chromosome arrival at the poles to ensure mitotic exit in time and cell survival (Woodruff et al., 2012). Our current knowledge of understanding spindle disassembly is mostly from the studies in budding yeast. The process of spindle disassembly in this organism is achieved by three pathways such as arrest of spindle elongation, depolymerization of interpolar MTs, and disengagement of spindle halves (Woodruff et al., 2010). These events are triggered and controlled by a signaling cascade termed the mitotic exit network (MEN) (Lee et al., 2001). Gene deletions and live-cell imaging assays have identified numerous MEN factors, such as kinases, phosphatases, molecular motors and molecular chaperones, participating in the regulation of spindle disassembly by inactivating spindle-stabilizing proteins and activating the spindle-destabilizing proteins (Ibarlucea-Benitez et al., 2018; Zimniak et al., 2009). The similar mechanisms have not been reported in other species, but it seems logically to speculate that mammalian cells have evolved more complicated machineries to control the disassembly. However, the spindle is disassembled in a very short time (within ~2 min), therefore, a simple switch-like mechanism is also expected likely at play in mammalian cells. Reduction-oxidation (redox) regulation could represent a candidate mechanism contributing to its highly dynamic and reversible properties perfectly matching the requirement of swift turnover of spindles.

In recent years, the intracellular redox state has been gaining increasing attention as crucial components involved in regulating cell proliferation (Schieber and Chandel, 2014; Wilson and González-Billault, 2015). Cells recruit diverse reactive oxygen species (ROS), including hydrogen peroxide (H_2_O_2_) produced during metabolism, to support proper cellular functions (Laurent et al., 2005; Rhee, 1999). A periodic redox oscillation during cell cycle was identified as one of crucial signals to modify many of key mitotic regulators, such as kinases and phosphatases (Jones, 2010; Menon and Goswami, 2006; Tu et al., 2005). Many genes involved in DNA replication or cell cycle progression have been shown to be expressed reliably in the reductive phase during a metabolic redox cycle (Tu et al., 2005). Accumulation of prooxidant levels in early G1 period of the cell cycle was demonstrated to be necessary for the transition from G1 to S phase in human cells (Menon et al., 2003). The depletion or over-expression of antioxidant enzymes, e.g. catalase, GPx4 and MnSOD, were shown to induce cell proliferation arrest (Brown et al., 1999; Sarsour et al., 2005; Wang et al., 2003).

More recently, MT dynamics is known to be coupled with the cellular redox homeostasis. Cysteine residues from α/β-tubulin can be oxidized by endogenous and exogenous oxidizing agents, resulting in its functional change on GTP binding or MT polymerization (Ludueña et al., 1985; Mellon and Rebhun, 1976). The treatment of peroxynitrite (ONOO-) on purified tubulin leads to glutathionylation of the thiol groups of tubulin monomers, thereby decreasing the polymerization of MTs *in vitro* (Landino et al., 2007). In addition, tubulin glutathionylation could be reversed by glutathione/glutathione reductase system (Landino et al., 2004a), suggesting that intracellular redox signal may modulate MT dynamics in a reversible manner. Besides, oxidation of cysteine residues on MT-associated proteins (MAPs) such as MAP2 and tau largely decreased MT polymerization *in vitro* (Landino et al., 2004b; Lewis et al., 1988), indicating that redox signals could regulate tubulin dynamics not only through direct interaction and modification, but also mediated by MAPs. Although these data have clearly pointed out the significance of redox regulation in cell proliferation and tubulin state, no any information is available concerning the direct relationship between redox signals and spindle dynamics.

Considering the complexity of their cysteine composition, we did not choose tubulin subunits but their associated proteins as candidates to test the possibility of redox driving spindle disassembly. In this study we identified that the spindle disassembly in human cells was controlled by redox modifications on cysteine residues in a tubulin-associated protein CRIPT (Cysteine-rich interactor of PDZ-domain protein). CRIPT is a highly conserved 101-residue protein containing four CXXC pairs expressed in animals, plants and single-cell eukaryotes (Figure 1). Human CRIPT shares 92 %, 70 %, 30 % sequence identity at the protein level with its homologs in frog, rice, and bread mold respectively. The feature having early evolutionary origin and remarkably high sequence conservation suggests it has a fundamental cellular function. We investigated CRIPT as an excellent candidate among tubulin-associated proteins who may have a redox-dependent impact on spindle disassembly relying on the lines of clues, including (1) The genomic analysis on patients with Primordial Dwarfism (PD) revealed that CRIPT is a disease associated gene and its depletion or mutation leads to fetal growth inhibition and stunted adults with microcephaly (Shaheen et al., 2014), indicating its roles in the cell proliferation control. (2) Mammalian CRIPT was previously demonstrated as a direct interactor of tubulin subunits (Niethammer et al., 1998; Passafaro et al., 1999), thus potentially participating in the regulation of spindle dynamics. (3) This protein has four extremely conserved CXXC pairs (Figure 1), while CXXC pair was identified as the most common type among disulfide bonds in bacterial proteins. (4) A naturally occurred single cysteine to tyrosine substitution (C3Y) in CRIPT gene was identified as a causative change in a PD patient (Leduc et al., 2016), suggesting the importance of this cysteine residue in cell division.

**Figure 1.**
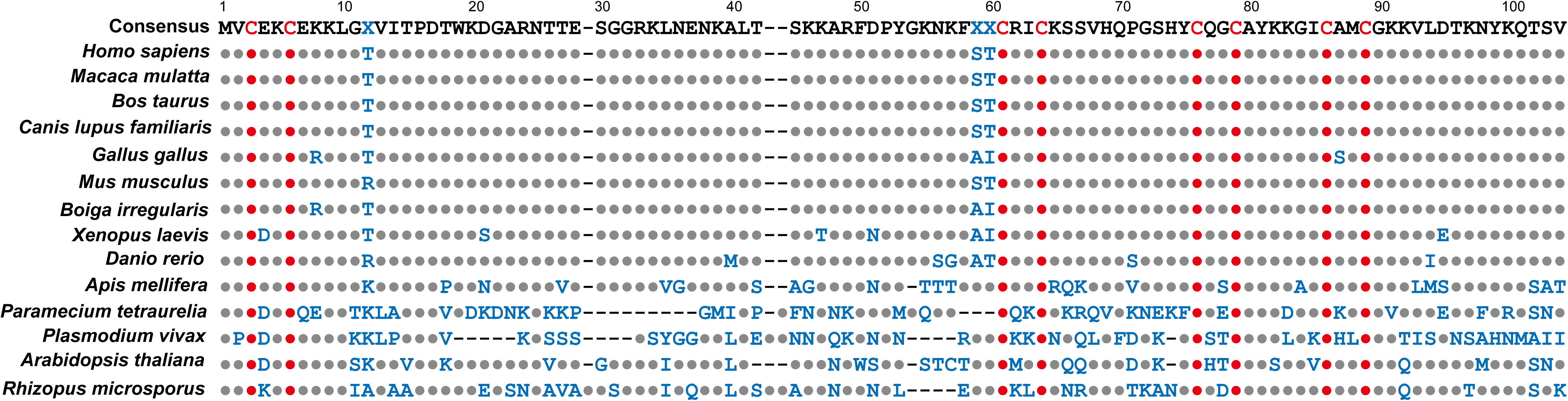
Sequence alignment of CRIPT in eukaryotes. The amino-acid sequences of CRIPT paralogs from diverse organisms, including *Rhizopus microspores*, *Arabidopsis thaliana*, *Plasmodium vivax*, *Paramecium tetraurelia*, *Apis mellifera*, *Danio rerio*, *Xenopus laevis*, *Boiga irregularis*, *Mus musculus*, *Gallus gallus*, *Canis lupus familiaris*, *Bos Taurus*, *Macaca mulatta* and *Homo sapiens*, were selected and aligned using Pfam 28.0 (www.pfam.xfam.org). The sequences are organized manually with the consensus amino-acid sequence on the top panel. The amino acids identical to consensus sequence are presented in grey solid circles, and the eight cysteines which are extremely conserved across all organisms are highlighted in red solid circles. The amino acids different from the consensus sequence are indicated in blue color.

In this study, we combined the cell imaging analysis and biochemical assays to define the key importance of CRIPT redox modifications on spindle disassembly in human cells. Apart from gaining the evidence to answer why CRIPT gene defects cause PD disease, our studies revealed that each of cysteine replacement in CRIPT protein completely impairs the disassembly of spindles to force them progress into an endless “tug of war” movement and redox modification on CRIPT protein is indispensable for MT depolymerization in two different mitotic phases. In addition, we had an unexpected finding on the cellular localization of CRIPT protein. In the previous reports, both endogenous and exogenous tagged (myc- and HA-tagged) CRIPTs were demonstrated to be functionally cytoplasmic proteins in neurons and interphase COS-7 cells (Niethammer et al., 1998; Passafaro et al., 1999; Zhang et al., 2017). Here, we determined CRIPT as a nuclear protein in neurons and other non-dividing cells.

## Results

### CRIPT is a spindle-associated nuclear protein

The genomic analysis connecting CRIPT defects to growth retardation predicts its participation in controlling the cell proliferation during development. To attest this prediction, we first knocked out this gene by CRISPR/Cas9 technology in cultured human cells and analyzed its effects on cell division. CRIPT depletion strongly increased the proportion of populations in mitotic metaphase compared with the parental cells (~15% versus ~2%). Besides, multipolar spindles and spindle misorientation were more frequently seen in the knockout cells (Figure 2A and B). These phenotypes support an essential role of CRIPT in the progression of cell division and in spindle organization. Next, to dissect its functions in greater depth, we expressed DsRed- and GFP-CRIPT in cultured human cells. Very unexpectedly, both fluorescence proteins were found to be restricted in nucleus and were highly concentrated in nucleoli with a good colocalization with nucleolar marker fibrillarin in interphase cells (Figure 2C-D and S1A). This localization of these fluorescence proteins is conflictive to the previous reports showing that endogenous or tagged CRIPT is a tubulin-associated cytoplasmic protein in cultured human COS7 cells and mouse neurons (Niethammer et al., 1998; Passafaro et al., 1999; Zhang et al., 2017). To exclude the possibility that DsRed and GFP tags caused the difference in subcellular distribution, the localization of endogenous CRIPT was studied in cultured human cells and cultured mouse neurons as well as mouse brain slices using both commercial and in-house anti-CRIPT antibodies. Consistently, endogenous CRIPT was stained in nucleus and found in the nuclear/nucleolar fraction (Figure 2E-F and S1A). After repeated examinations of different tagged and endogenous CRIPT in tens of different cell lines, we defined CRIPT as a nuclear protein. Further, we found that CRIPT localizes along the spindles throughout cell division and concentrates in spindle poles (Figure 2G and S1B). Therefore, CRIPT is also a spindle associated protein during mitosis.

**Figure 2.**
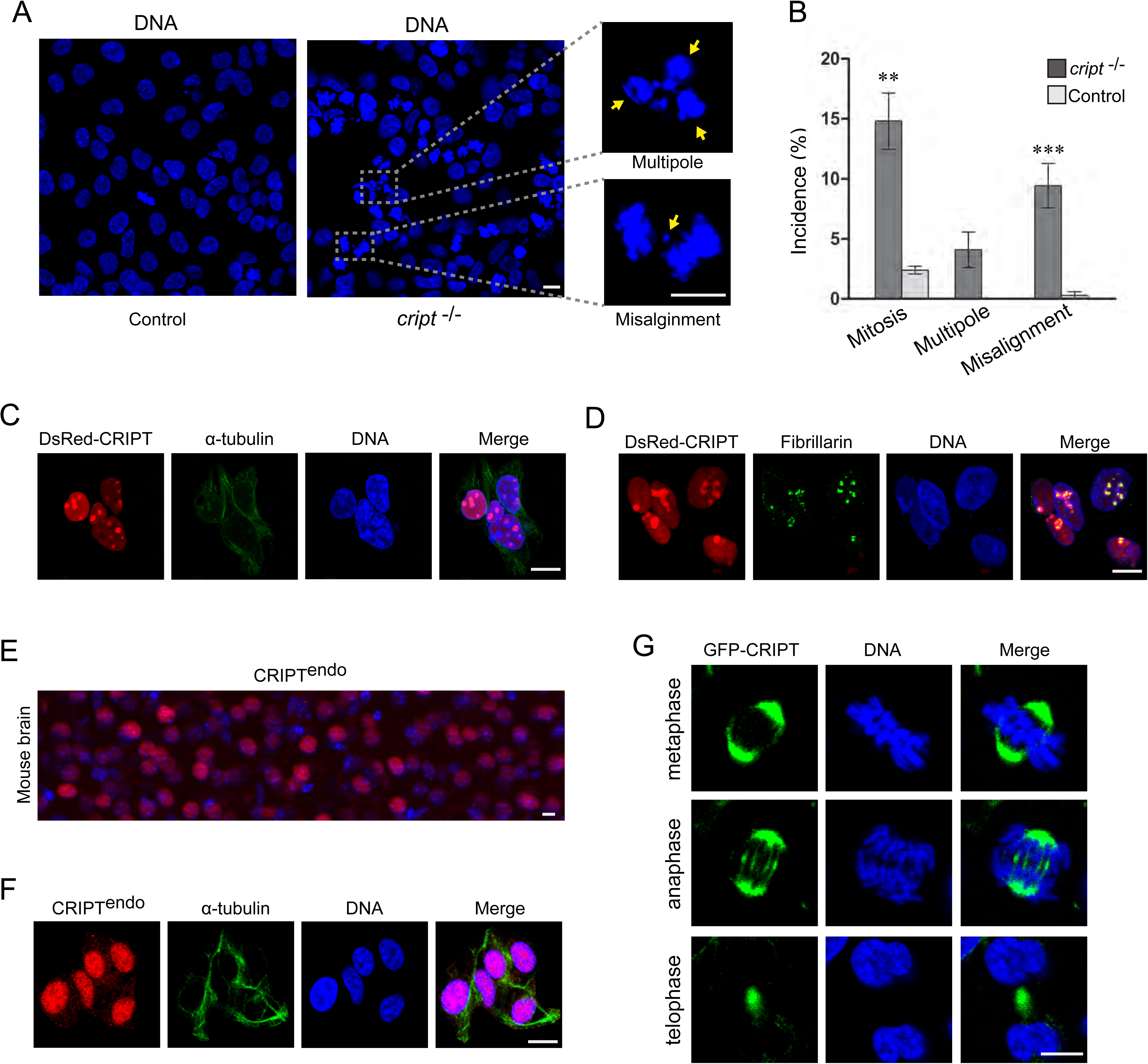
DNA morphology of CRIPT knockout cells and subcellular localization of the native and tagged CRIPT proteins. **(A)** CRIPT knockout (*cript* ^−/−^) HeLa cells are frequently visualized in mitosis with multipolar spindles or lagging chromosomes. Arrows point to multiple spindle poles or lagging chromosomes. Untreated HeLa cells are used as normal control. **(B)** Schematic representation of *cript* ^−/−^ cells displaying mitotic defects. Data are presented as mean ± s.e.m. and unpaired t-test was used for data analysis. *^**^P* < 0.01, *^***^P* < 0.001. **(C)** HeLa cells expressing DsRed-CRIPT (red) are immune-stained for α-tubulin (green). **(D)** HeLa cells expressing DsRed-CRIPT (red) are immune-stained for fibrillarin (green) to show their colocalization. **(E)** The merged imaging showing endogenous CRIPT (red) colocalizing with DNA (blue) in a mouse brain slide. **(F)** HeLa cells are immune-stained for endogenous CRIPT (red) and α-tubulin (green). **(G)** HeLa cells expressing GFP-CRIPT in three mitotic phases are shown. DNAs in **A** and **C-G** are visualized by counterstaining with DAPI (blue). Scale bars, 10 µm.

### CRIPT functions as a determining factor in spindle disassembly

C3Y mutation in CRIPT was reported to be associated with symptoms of short stature and microcephaly in a female, suggesting that this cysteine residue might be important for its physiological function. To test this idea, we first expressed DsRed-CRIPT^C3Y^ and -CRIPT^WT^ in HeLa cells and compared their cellular distribution and effects in cell morphology and proliferation. Both fluorescence proteins were restricted in nucleus at 24 hours after cell transfection. Later, different from DsRed-CRIPT^WT^ who did not alter normal cell division, DsRed-CRIPT^C3Y^ gradually increased two types of cell populations (Figure 3A-D). These cells were found to be arrested in mitosis based on α-tubulin and DNA staining. Type A cells were characterized by DsRed-CRIPT^C3Y^ localizing along the spindles with two bright spots at poles (we named them as “bi-pole ring” spindles) (Figure 3B), indicating a defect of spindle poleward movements. Type B cells were sorted by DsRed-CRIPT^C3Y^ decorating a messy fiber-like structure (Figure 3C). Through careful analysis, we diagnosed that this structure might be a remnant from undissolved spindles. At 48 hours after transfections, type A and B cells constituted 34% and 22% populations in DsRed-CRIPT^C3Y^ expressed cells, respectively (Figure 3D). These morphological data revealed that C3Y mutation in CRIPT exerted an inhibitory effect on cell cycle progression, and most likely through interfering with spindle disassembly.

**Figure 3.**
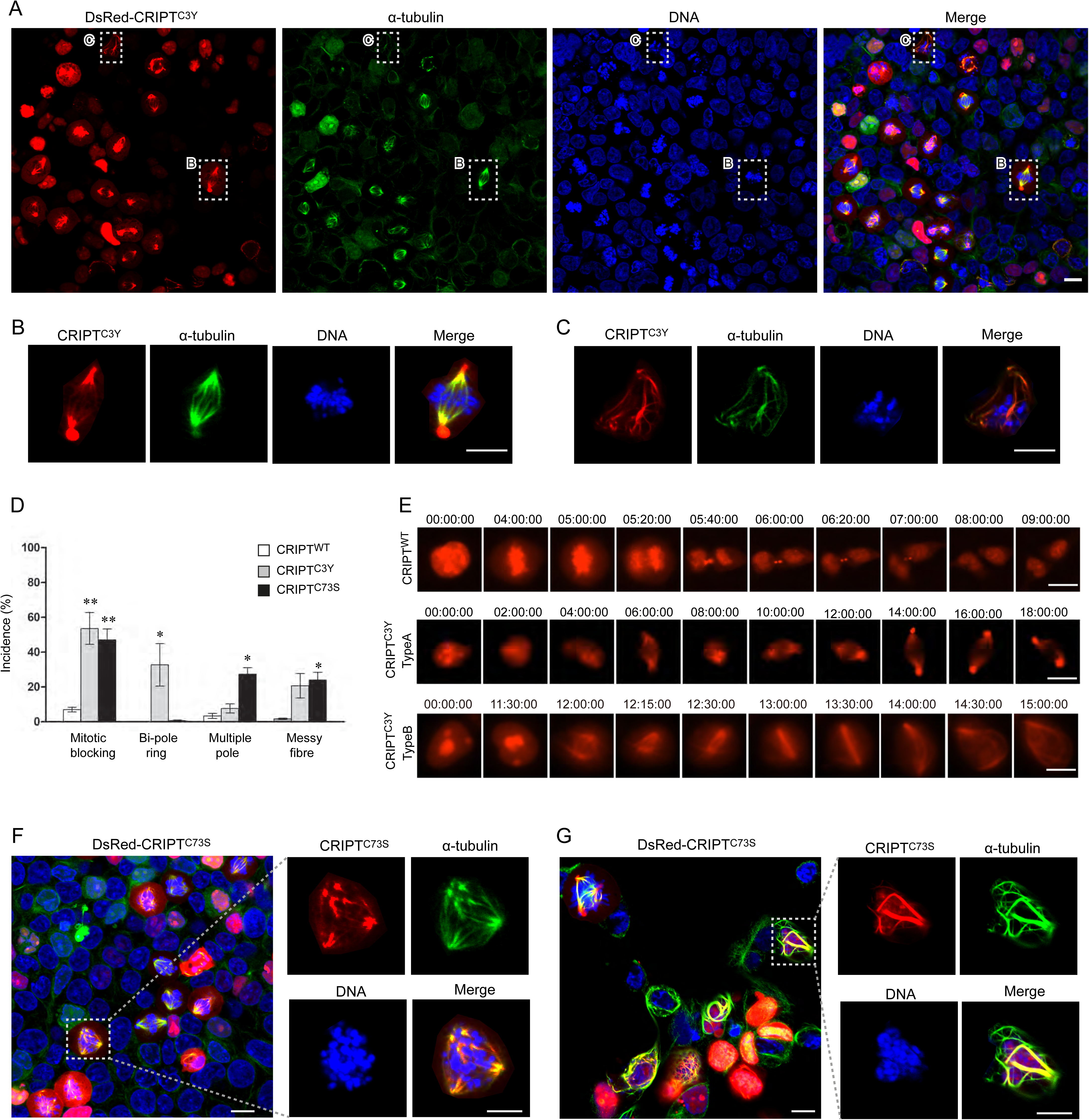
The phenotypes and time-lapse imaging of HeLa cells expressing CRIPT mutants. **(A)** HeLa cells expressing DsRed-CRIPT^C3Y^ (red) were immune-stained for α-tubulin (green) and DNA was counterstained with DAPI. **(B)** Inset from **A** showing the high-magnification (5×) image of a cell having Type A spindle. **(C)** Inset from **A** showing the high-magnification (5×) image of a cell having Type B spindle. **(D)** Schematic representation of cells expressing CRIPT mutants with diverse spindle configuration. Abnormal spindle count in cells expressing CRIPT^WT^ was used as control. The data was collected from three fields of each transfection and the error bars represent mean ± s.e.m. *^*^P* < 0.05; *^**^P* < 0.01. **(E)** HeLa cells transfected with DsRed-CRIPT^WT^ (top panel) and -CRIPT^C3Y^ (middle and bottom panel) were monitored using live cell microscopy for 24 hours. Frames taken at the indicated time points (h : m : s) were shown. **(F)** HeLa cells expressing DsRed-CRIPT^C73S^ (red) were immune-stained for α-tubulin (green) and DNA was counterstained with DAPI. **(G)** Inset from F showing the high-magnification (5×) image of a cell with multiple-pole spindles. **(H)** Inset from F showing the high-magnification (5×) image of a cell with disordered fiber-like spindle.

To further and more clearly define the roles of CRIPT in spindle disassembly, we tracked the spindle dynamics using time-lapse fluorescence microscopy in cells expressing DsRed-CRIPT^C3Y^ or DsRed-CRIPT^WT^. The spindles were visualized by fluorescence CRIPT proteins rather than most often used tubulin subunits. As shown above, CRIPT redistributes into the cytosol from nucleus at the onset of mitosis and fully decorates the spindles during mitosis. Thus, we considered CRIPT as a better marker to monitor spindle dynamics together with the cell cycle progression relative to tubulin subunits. Cells expressing CRIPT^WT^ were able to accomplish their division within 2-3 hours (Figure 3E top panel and Video 1). Once entering the mitosis, cells rapidly progressed into anaphase within 20 min. Following the complete separation of two spindle halves in telophase, the entire spindles are thoroughly dissolved within 2 min. Then, CRIPT relocated and concentrated in midzone where persisting for more than 1 hour till the end of cytokinesis (Figure 3E top panel and Video 1). In the cells expressing CRIPT^C3Y^, we recorded two types of cells with distinct spindle dynamics in compatible with two cell populations described above (Type A and B cells). In type A cells (Figure 3E middle panel and Video 2), the metaphase spindles were regular as seen in CRIPT^WT^ cells, but they were hyperstable and lasted more than 8 hours without any noticeable changes in length of each spindle and in distance between two spindle halves. Thereafter, some fluorescence accumulated at the centrosomes to form the “bi-pole ring” spindles and cells stayed in this state until the cell death. These images revealed that the hyperstable spindles resulted from failure to separate spindle halves due to defects in poleward movements, and explained well why the cells with “bi-pole ring” spindles are mostly captured under laser confocal microscopy. In type B cells (Figure 3E bottom panel and Video 3), the metaphase spindles were also morphologically normal and persisting, however, they were highly dynamic. Both spindle halves displayed a continuous “tug of war” movement for 3-4 hours, seeming they were “struggling” to break apart. Finally, when completely separated, they were not dissolved but rapidly gathered to form a long-lasting fiber-like structure, constituting a spindle remnant. Obviously, these live cell images from type B cells provide a clear proof that a fiber-like structure in CRIPT^C3Y^ cells is a spindle remnant as a consequence of defects in spindle disassembly. Altogether, these data from real-time live imaging confirmed that CRIPT^C3Y^ led to its phenotypes through blocking the spindle poleward movements in anaphase or/and spindle dissolution in telophase. It is generally believed that in addition to be responsible for spindle dissolution in telophase, MT disassembling is a major mechanism for the spindle poleward movements in anaphase. Thus, CRIPT^C3Y^ undoubtedly had a negative effect on MT disassembly. Also, this finding provides insight into a selected mutation in CRIPT associated diseases in human with growth retardation and microcephaly.

Next, considering that the cysteine residue is subject to be modified to form intra/inter-molecular disulfide and eight cysteines in CRIPT are extremely conserved, we were interested in exploring the involvement of other seven cysteines in determining the fate of spindles. We systematically observed the behaviors of a series of CRIPT mutants with a single or combined cysteine substitutions by a serine or alanine residue. Surprisingly, all these mutants led to phenotypes of two populations (Type A and B cells) containing long-lasting spindles or spindle remnants in transfected HeLa cells as seen in CRIPT^C3Y^ expressed cells. This result not only highlights the importance of these cysteine residues for CRIPT function, but also suggests that eight cysteines might work as a whole. However, after carefully examining the spindle morphology in type A cells, we found that eight cysteines were also differentiated functionally as summarized in Table 1. Mutants (CRIPT^S4^) with substitutions to any of first four cysteines (such as C3S, C6S, C58S, C61S, C3/6S and C58/61S) generated typical “bi-pole ring” spindles as seen in C3Y expressed cells, whereas mutants (CRIPT^L4^) with substitutions to any of last four cysteines (such as C73S, C76S, C83S, C86S, C73/76S and C83/86S) induced multiple-pole spindles (Figure 3D, F and G), which indicates the defects in organizing MT center. All these mutants generated similar two cell populations in other ten tested cell lines such as HEK293, HepG2, COS7, OV90. Therefore, eight cysteines in CRIPT are functionally hierarchical. Their roles in disassembling the spindles are uniform, but are different and grouped in organizing MT center. To exclude the possibility that mutant-induced phenotypes might result from artificial or detrimental effects due to residue replacements, we also tested other three mutants (W19S, N37S, H71A) created by a random residue replacement with a serine/alanine residue. These mutants all behaved like CRIPT^WT^ exclusively present in nucleus in interphase cells, and none of them induced long-lasting spindles. This result further confirmed the importance and specificity of these cysteines. Altogether, we identified CRIPT to be a crucial factor in determining spindle disassembly, and revealed the importance of its cysteine residues in the course of performing this task.

**Table 1.**
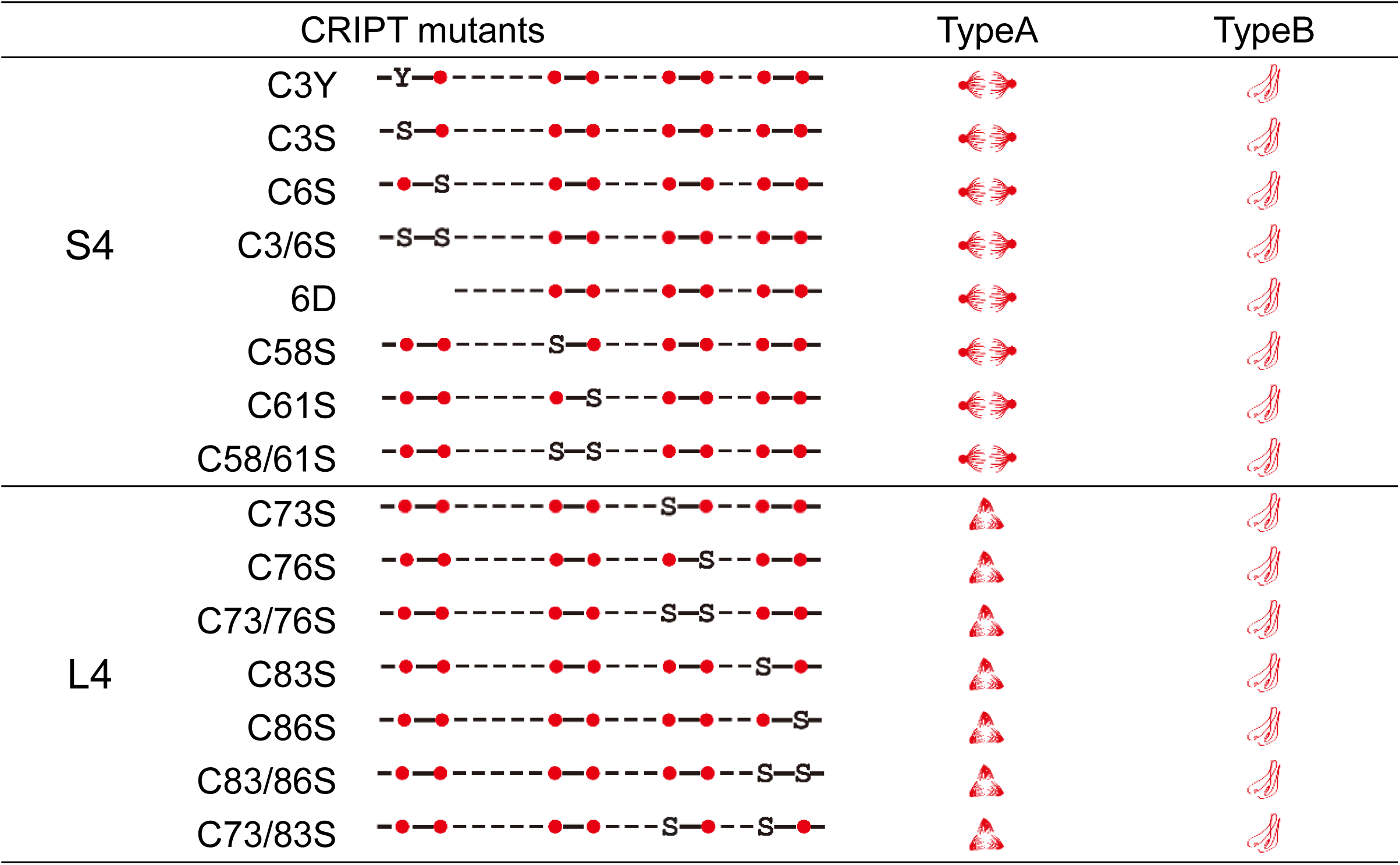
Spindle configuration of the cells expressing CRIPT mutants.

### CRIPT is a cysteine-dependent tubulin binding protein

The imaging results that overexpressed CRIPT and its derivatives had a restrictive distribution on mitotic spindle urged us to explore the possibility of the physical interaction between CRIPT and microtubule proteins. The association of CRIPT with α-tubulin has been demonstrated in previous studies although their results on CRIPT localization are conflictive to ours. In our co-immunoprecipitation assays, CRIPT and α-tubulin were brought down each other from the lysates of transfected mitotic cells (Figure 4A), supporting that they are in a same molecular complex in mitotic cells. To confirm their direct interactions, GST pull down assays were performed using purified bacteria expressed his-tagged CRIPT and GST-tagged tubulin subunits. As shown in Figure 4B, his-tagged CRIPT were eluted down with each of three tubulin subunits, indicating a direct association of CRIPT with tubulin. These results strongly recommend CRIPT as a novel tubulin binding protein. Many MT-binding proteins have a common CAP-Gly domain, forming a positive charged groove on the molecular surface to interact with the C-terminal acidic tails of tubulin subunits (Figure 4C). The core sequence of microtubule recognition in a CAP-Gly domain is a GKNDG motif with a critical importance of the Lys residue whose alanine substitution leads to a significant reduction of tubulin-binding affinity (Mishima et al., 2007). CRIPT has a nearly same amino acid sequence of GKNKF at position 51 to 55, accordingly, this sequence was investigated as a potential α-tubulin binding motif. Bacteria expressed his-tagged CRIPT^ΔGKN^ and CRIPT^K52A^ were examined along with GST-tagged α-tubulin in pull down assays to determine the importance of this sequence. As expected, both mutations resulted in a significant reduction of the α-tubulin binding affinity (Figure 4D). This result not only helped us identify an interacting motif, but also strengths our conclusion that CRIPT is a tubulin interacting protein.

**Figure 4.**
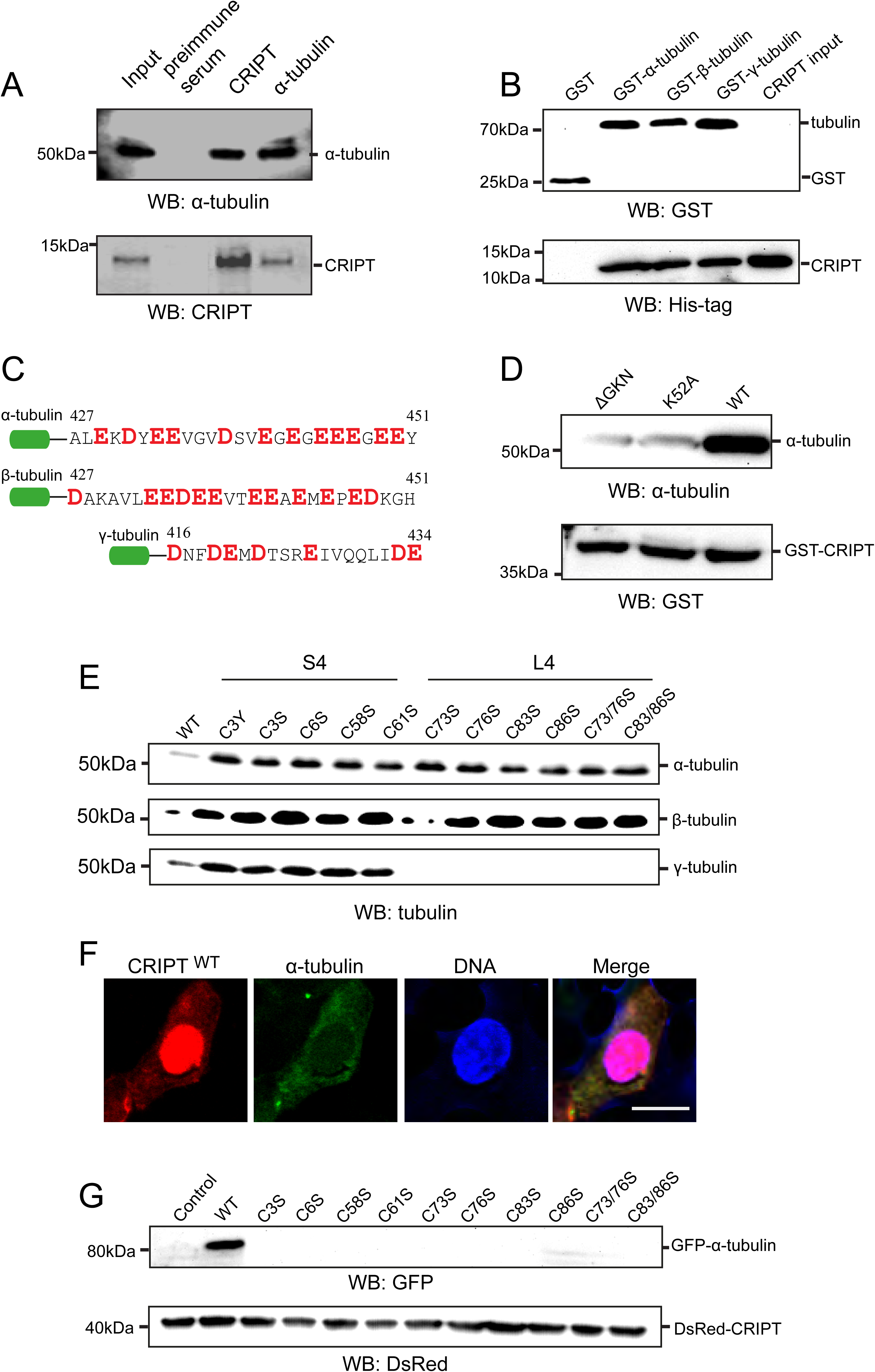
Cysteine residues are necessary for microtubule binding of CRIPT. (**A**) The interaction between CRIPT and α-tubulin *in vivo*. The lysates from mitotic HeLa cell were immunoprecipitated with an anti-CRIPT antibody followed by detection of α-tubulin. The same blot was probed with a polyclonal CRIPT antibody. Pre-immune serum was used as the negative control in the blot. (**B**) *In vitro* binding assay with GST-tagged tubulin subunits and purified His-tagged CRIPT. The blot was probed using monoclonal anti-His and anti-GST antibody, respectively. (**C**) The C-terminal acidic tails of three tubulin subunits. Aspartic acid (D) and Glutamic acid (E) were highlighted in red and bold font. (**D**) Pull-down assays of α-tubulin with GST-CRIPT^WT^ and -CRIPT mutants. CRIPT^WT^ and CRIPT with mutagenesis on lysine (CRIPT^K52A^) or depletion of “GKN” motif (CRIPT^ΔGKN^) were immobilized on the resin and tubulin was used as an input. (**E**) Coimmunoprecipitation assays on CRIPT protein with cysteine substitution at different positions with endogenous α, β, γ-tubulin subunits. (**F**) DsRed-CRIPT cytoplasm distribution when co-expressed with α-tubulin in HeLa cells. (**G**) Coimmunoprecipitation assays on DsRed-CRIPT cysteine-substituted mutants with GFP-α-tubulin in interphase HeLa cells. Cell lysates from transfected DsRed-CRIPT alone were used as control.

To understand the reason why cysteine mutated CRIPTs were tethered to and fix the mitotic spindle, we focused on investigating the roles of cysteine residues on its associations with tubulin subunits. A series of CRIPT mutants with a single or combined cysteine substitutions were transfected into HeLa cells and their associations with endogenous tubulin subunits were detected by co-immunoprecipitation assays at 48hr after transfections. Consistent with their accumulation on the mitotic spindle, all of these cysteine-mutated CRIPT relative to its wild-type (WT) protein brought down more than two-fold α and β-tubulin from the mitotic cells (Figure 4E), indicating that these cysteine mutations endow the derivatives with an enhanced binding affinity to mitotic α and β-tubulin. Meanwhile, no significant difference on the binding affinity among these derivatives suggests that eight cysteine residues in CRIPT could be functionally uniformed or as a whole in the formation of partners with α /β-tubulin. However, the roles of eight cysteine residues were found to be different and grouped when the association of CRIPT with γ-tubulin was examined. As shown in Figure 4E, CRIPT^S4^ displayed an enhanced binding affinity to γ-tubulin compared to the CRIPT^WT^ protein, whereas CRIPT^L4^ completely lost the capability of γ-tubulin binding. These results may provide a direct evidence of the involvement of cysteine residues in the association of CRIPT with mitotic spindle. However, with regard to the fact that these associations were examined in mitotic cells, the change of binding affinity could only occur on tubulin oligomers (mitotic microtubules), but not on the individual tubulin protein. To be able to decipher the roles of CRIPT cysteines in the associations with individual tubulin protein, α, β, γ tubulin subunits were respectively co-transfected into cells with CRIPT^WT^ and CRIPT mutants, and their associations were studied in the interphase cells. Different from the endogenous and the single overexpressed proteins being exclusively confined to the nucleus, there are substantial CRIPT^WT^ distributed into the cytoplasm in the interphase cells when each of three GFP-tagged tubulin were co-overexpressed in the same cells. Figure 4F is a sample showing an interphase HeLa cell expressing GFP-tagged α-tubulin and DsRed-CRIPT. This phenomenon that CRIPT protein can be retained in the cytosol by co-overexpressed tubulin proteins represents a strong evidence of their direct interactions in cells. Conveniently, we took use of this phenomenon to diagnose the effects of cysteine mutations on CRIPT-tubulin interactions via the direct observations in the cotransfected cells. In the following transfection assays, a series of cysteine-substituted DsRed-CRIPT with GFP-tagged tubulin subunits were co-transfected into cells, and cells were arrested in the interphase by serum withdraw at 24hr after transfection. Our observations at 48hr after transfection revealed that except WT protein, none of cysteine-substituted CRIPT mutants appeared in cytosol. No cytoplasmic distribution of these mutants should mean a loss of CRIPT association with tubulin subunits in cells after its cysteine replacements. This prediction was confirmed by the co-immunoprecipitation assays performed from the interphase cells, in which the interactions with tubulin proteins were detected only for CRIPT^WT^ but not for any of cysteine-substituted mutants (Figure 4G). Despite the fact that the behaviors of cysteine substituted mutants in mitotic and interphase cells were found to be seemingly discrepant, these results have verified their critical roles in the formation of CRIPT-tubulin partners. On the other hand, this discrepancy may reflect the variable roles of CRIPT cysteines in different cellular environments.

### CRIPT CXXC pairs are modified by redox signals during mitosis

The above mutagenesis assays in the transfected cells revealed that each of eight cysteine resides in CRIPT is of functionally important, in agreement with our assumption of its involvement in redox regulation of cell cycle progression. These cysteines may natively constitute key structural elements, alternatively they may undergo certain modifications which are required to fulfil its cellular functions. Considering that a serine residue has a good structural match with a cysteine residue, the fact that the serine substitution in every position caused a completely functional loss argued against a naturally structural role. Importantly, eight cysteines in CRIPT are present in four typical CXXC pairs, which is the most common type among disulfide bonds. To gain a mechanistic insight into these cysteine pairs, we focused our studies on investigating whether they were involved in the formation of intramolecular/intermolecular disulfide bonds. For this purpose, we first assayed redox state of these cysteines in the cultured cells.

Based on the clues of functional studies, it is plausible to infer that these cysteines may have differential redox states in a certain environment. To simplify our assays, we wanted to examine 8 cysteines in a whole rather than individually in the first step, therefore, only CRIPT^WT^ and cysteine-lacking mutant (CRIPT^8C-S^) were included in the preliminary experiments. His-tagged CRIPT^WT^ and CRIPT^8C-S^ were expressed in HeLa cells and mitotic cell lysates were directly examined by anti-His immunobloting. The results showed that CRIPT^WT^ and CRIPT^8C-S^ appeared indistinguishably as a single species with an expected 14KD size under both non-reducing and reducing conditions (Figure 5A). No additional higher species under a non-reducing condition indicates no stable intermolecular disulfides in CRIPT being formed up to a detected level in cells. However, no protein band shift under different redox conditions cannot rule out the existence of intramolecular disulfide bonding, although intramolecular disulfides can cause slightly faster migration in some proteins under a non-reducing condition. To further explore the intramolecular disulfide bonds, the purified His-tag proteins were examined by Coomassie-blue staining and anti-CRIPT immunoblots (Figure 5B and C). Surprisingly, CRIPT^WT^ migrated slightly slower under a non-reducing condition than under a reducing condition, whereas CRIPT^8C-S^ did not show any migration difference under both conditions, suggesting the possible oxidative modifications such as intramolecular disulfide bonds forming on cysteine residues. When proteins were loaded up to 2µg in one gel well, the additional higher species were visualized clearly by Coomassie-blue staining in the non-reducing sample of CRIPT^WT^ proteins but not in CRIPT^8C-S^ protein sample, and those higher species were later revealed to be the homodimer or homoligomer by anti-CRIPT immunoblot, indicating the formation of intermolecular disulfide bonds on cysteine residues. To confirm the existence of disulfide bonds in purified proteins, the biotin switch method was used. CRIPT^WT^ but not CRIPT^8C-S^ was labeled by MTSEA-biotinylation (Figure 5D), conferring the presence of cysteine-dependent disulfide bonds in CRIPT^WT^ protein. The results demonstrated that CRIPT has redox reactive cysteines, consistent with the mutagenesis assays in the transfected cells.

**Figure 5.**
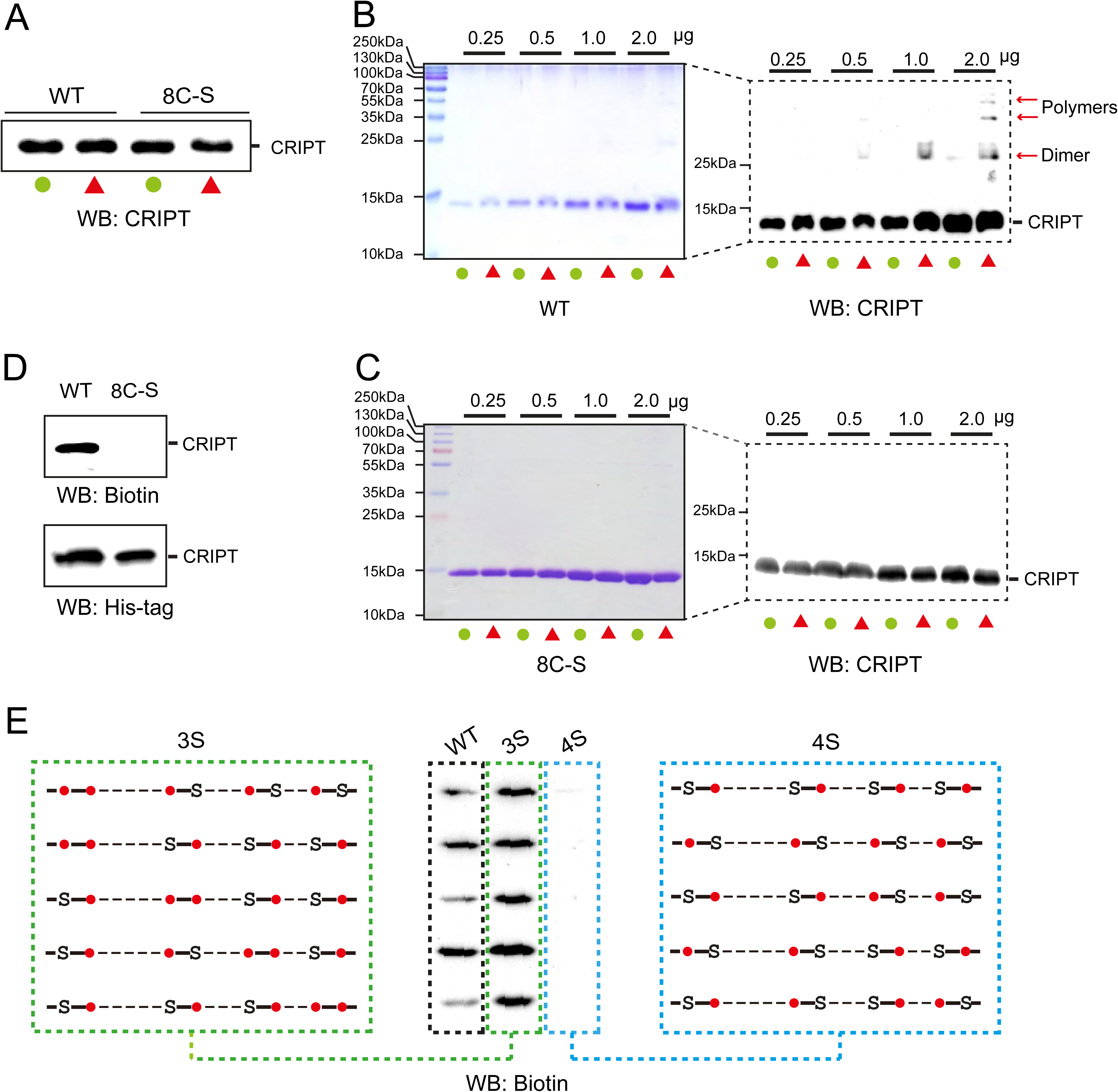
Disulfide bonds in CRIPT CXXC pairs are formed during mitosis. (**A**) His-tagged CRIPT was expressed in HeLa cells and examined by anti-His blotting under both non-reducing and reducing conditions. (**B-C**) Purified His-tagged CRIPT and CRIPT^8S^ denatured in reducing and non-reducing conditions were separated on native PAGE and examined by both Coomassie blue staining and anti-CRIPT immunoblots. Non-reducing and reducing condition treatment were indicated by green circle and red triangle, respectively. (**D**) CRIPT and 8C-S variant were labeled with *biotin-linked* cysteine-specific reagent (*MTSEA-*biotin) and examined by biotin-streptavidin system. (**E**) CRIPT variants with combined cysteine substitutions were labeled with *MTSEA-*biotin and examined by biotin-streptavidin system. The schematic representation of diverse combination of cysteine substitutions were shown, cysteine residues and the substitutions were indicated as red circle and amino-acid codes, respectively.

Next, to determine the pattern of disulfide forming, a series of CRIPT variants with combined cysteine substitutions were examined using the same biotin switch method (Figure 5E). The collective variants having one of CXXC pairs intact (designated as CRIPT^3S^) were all labeled by MTSEA-biotinylation even stronger that CRIPT^WT^ protein, while the collective variants have a substitution in each of CXXC pairs (designated as CRIPT^4S^) were not labeled, indicating that disulfide bonds were formed within CXXC pairs. The stronger labeling of CRIPT^3S^ variants than CRIPT^WT^ protein implies that the formed disulfide bonds in CRIPT^WT^ protein are more mobile or short-lived. Furthermore, the existence of a disulfide within a CXXC pair was found to be a prerequisite for the formation of intermolecular disulfides *in vitro* because no dimers/oligomers was detected in CRIPT^4S^ protein.

### The first two CXXC pairs of CRIPT participate in a thiol/disulfide exchange with tubulin

The emerging evidence has suggested that the organization of the tubulin cytoskeleton seems to be tightly associated with the cellular redox status. The highly dynamic organization of mitotic spindle may be achieved as a balance between redox-regulated effects on tubulin polymerization and depolymerization. The above findings that CRIPT cysteines are under a very mobile redox state in mitotic cells and that redox-deficient mutants abolished the dynamic change of mitotic microtubules so as stabilizing them at a polymerization state have indicated the potential redox interactions between CRIPT and tubulin subunits during mitosis. Therefore, we speculated that CXXC pairs in CRIPT could react with tubulin cysteines by a thiol/disulfide exchange mechanism. To test this hypothesis, we examined whether CRIPT could enhance tubulin oxidation by H_2_O_2_ to form disulfide bonds *in vitro*. Purified tubulin proteins alone or with CRIPT were treated with H_2_O_2_, and detected by anti-α-tubulin immunoblotting under nonreducing conditions. As shown in Figure 6A, the higher molecular weight species representing oxidized α-tubulin oligomers appeared in the sample with CRIPT addition, whereas only dimers were found in the presence of tubulin alone, which is consistent with the speculation that oxidation of tubulin cysteines by H_2_O_2_ can be facilitated by CRIPT. This effect of enhanced oxidation on tubulin may be only involved in first two CXXC pairs because the mutant CRIPT^L4^ reaction is similar to CRIPT^WT^. To further confirm the possible effects of CRIPT on H_2_O_2_ oxidation of tubulin thiols and determine which CRIPT cysteine pairs are responsible for these redox interactions, IAF labeling was used to monitor changes to tubulin cysteines in the presence of CRIPT and the derivatives. Because IAF only reacts with reduced cysteine, the intensity of IAF labeling directly indicates the number of the reduced cysteines in a molecule. CRIPT^WT^ was observed to significantly decrease the intensity of IAF-labeled α- and β-tubulin (Figure 6B), supporting it as a redox regulator of tubulin subunits. To understand the role of individual CXXC pairs, several CRIPT mutants were included in the same IAF labeling assays. CRIPT^L4^ was found to have an almost equal effect as CRIPT^WT^ protein to decrease tubulin labeling (Figure 6B), denying last four cysteines as the main participants in transducing CRIPT redox response to tubulins In contrast, CRIPT^S4^ completely lost the enhanced effects on tubulin oxidation (Figure 6B), implying the necessity of the first two CXXC pairs in the process of redox modifications from CRIPT to tubulin subunits. Collectively, we proposed that the first two cysteine pairs in CRIPT are mainly responsible for the redox interactions with tubulins possibly via a thiol/disulfide exchange mechanism.

**Figure 6.**
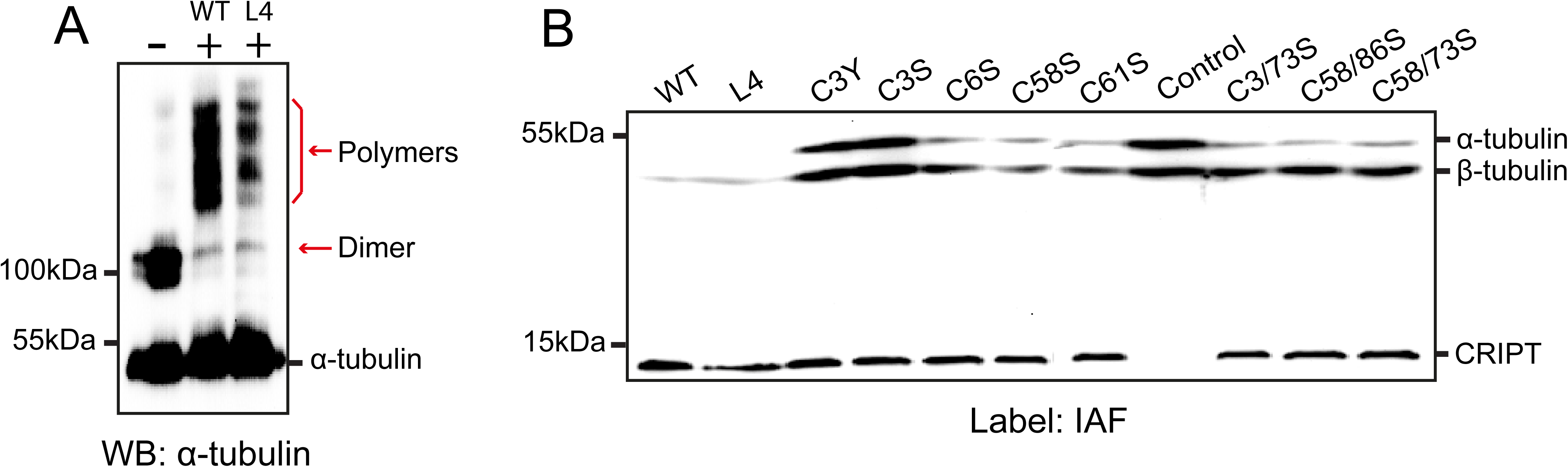
CRIPT enhances the oxidation of tubulin subunits by H_2_O_2_ *in vitro*. **(A)** Purified tubulin protein incubated with and without CRIPT were treated with H_2_O_2_ and examined by anti-α-tubulin immunoblotting. Arrows indicate oxidized α-tubulin. **(B)** Tubulin subunits in the presence of H_2_O_2_-treated CRIPT and CRIPT variants were labeled with IAF to monitor the reduced cysteines on these proteins.

### TBCB may facilitate CRIPT-mediated spindle disassembly

The above findings have identified CRIPT and related redox response as a critical component for spindle disassembly in human cells. However, disassembling the spindle is a highly complicated event, which is coordinately controlled in yeast by functionally distinctive protein complexes including motor proteins and molecular chaperones (Ibarlucea-Benitez et al., 2018; Zimniak et al., 2009). We suspected that CRIPT may function in a dependent protein complex to assist the spindle disassembly. Accordingly, we set out to identify and characterize possible CRIPT-associated disassembly factors in human cells. CRIPT complexes were isolated from the mitotic HeLa cells, and mass spectrometry (MS) analysis revealed some clues of CRIPT interactors, including β-actin, TPM3, ACRT2, CAPZB, TBCB and APRC1B (Figure 7A). Direct associations of CRIPT with most of these proteins were validated by GST pulldown assays (Figure 7B), although consequences of these associations are still unclear. Among these potentially functional partners, TBCB is of particular interest to us because it is a tubulin-specific chaperone mainly involved in α/β tubulin heterodimer dissociation, thus was further investigated with CRIPT in this study. Cells with CRISPR-cas9 mediated depletion of TBCB had a higher rate of mitotic population (Figure 7C), a phenotype closely similar to that in CRIPT null cells, implying the behavior relevance between CRIPT and TBCB. In transfected cells, fluorescent labeled TBCB was found to be distributed evenly in interphase cells, but concentrated at the metaphase spindle with a good colocalization with CRIPT (Figure 7D), morphologically providing a support for its CRIPT association. This colocalization on spindles is also independent on redox-competence of CRIPT as shown in Figure 7E. Based on these results, we speculate that TBCB could work together with CRIPT to prompt the spindle disassembly.

**Figure 7.**
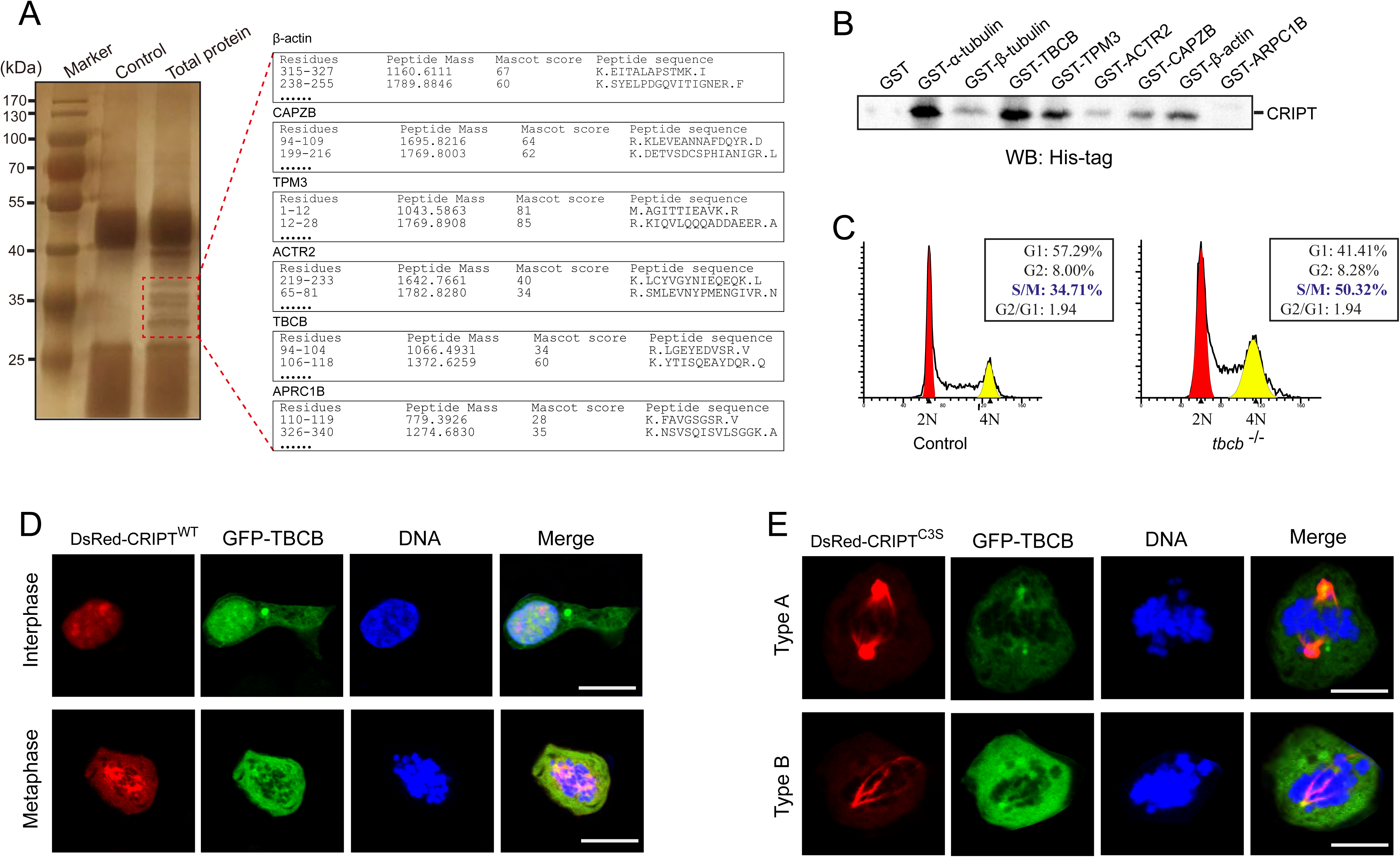
CRIPT colocalizes and interacts with TBCB. (**A**) Identification of TBCB as a CRIPT-associated protein by coimmunoprecipitation and MS assays. Inset is the list of matched peptides from MS analysis. (**B**) Validation of the interaction between CRIPT and MS-identified proteins by GST pulldown assays. (**C**) DNA contents of *tbcb*^−/−^ cells were analyzed by flow cytometry. The normal HeLa cells were used as control. (**D-E**) HeLa cells expressing DsRed-CRIPT^WT^ or -CRIPT^C3S^ and GFP-TBCB were examined using confocal microscopy to visualize their spatial relationship during mitosis. DNA was counterstained with DAPI. Scale bar, 10 μm.

## Discussion

### Spindle disassembly in human cells is controlled by CRIPT-bearing redox signals

The process of mitotic spindle disassembly is a complicated and poorly understood event. There are at least three pathways proposed to control spindle disassembly (Woodruff et al., 2010) in yeast cells, but none of them has been identified in mammalian cells. It is also unknown whether mammalian cells utilize the same or distinct mechanisms to manage their spindle disassembly in two different phases (Asbury, 2017). In this study, CRIPT was found to control the spindle disassembly via a thiol/disulfide exchange mechanism. By systematic analysis of a series of cysteine-mutated CRIPT variants, we obtained some unexpected findings to uncover the mystery how mammalian cells disassemble their spindles during mitosis. First of all, each of redox-deficient CRIPT mutations can completely block spindle disassembly, implying CRIPT-bearing redox signals as an essential mechanism. This is a totally unexpected finding because the intracellular redox signaling has never been appreciated to play such a crucial role in spindle disassembly. Secondly, each of redox-deficient CRIPT mutations was found to yield two cell populations with either spindle remnants or hyperstable metaphase spindles. The phenotype of hyperstable metaphase spindle indicates the failure of spindle separation due to the defect of spindle poleward movement in anaphase, while existence of spindle remnants is a consequence of spindle disassembly failure in telophase. The presence of mixed cell populations suggests that the disassembly of spindle ends during anaphase and the entire spindle disassembly during telophase may share a common redox-dependent mechanism. Obviously, this mechanism identified in human cells is completely distinct to those in yeast which are mainly carried out by protein phosphorylation and proteasome-mediated protein degradation (Ibarlucea-Benitez et al., 2018; Zimniak et al., 2009). However, this mechanism may not be unique in human cells given the high conservation of CRIPT protein and particularly of its cysteine pairs in most of species. On the other hand, human cells could have compensatory mechanisms to overcome the redox dependence of MT disassembly on the spindle ends to finish poleward movement, because co-existence of two cell populations indicates that some cells have ignored the defect of CRIPT-bearing redox signals in anaphase to enter telophase.

Although the detailed mechanism of spindle disassembly is waiting to address, our data has demonstrated that CRIPT resides at a major site where the disassembling signals are transferred to MTs. Given that most of cysteine residues in tubulin heterodimer are buried inside, an allosteric assistance on tubulin from the molecule chaperones is expected when transferring redox signals from CRIPT to MTs. A tubulin-specific chaperone TBCB has been shown to directly interact with CRIPT, therefore, CRIPT-bearing redox signaling could pass on tubulin under the help of TBCB to fulfill the final event---spindle disassembly. Here we outline a simple model in Figure 8 to explain how the redox signaling controls spindle disassembly in human cells, in which CRIPT senses and reacts with intracellular oxidants, then transfers redox responses and oxidize MT cysteines via a thiol/disulfide exchange under the allosteric assistance of TBCB up to a threshold number causing tubulin deploymerization.

**Figure 8.**
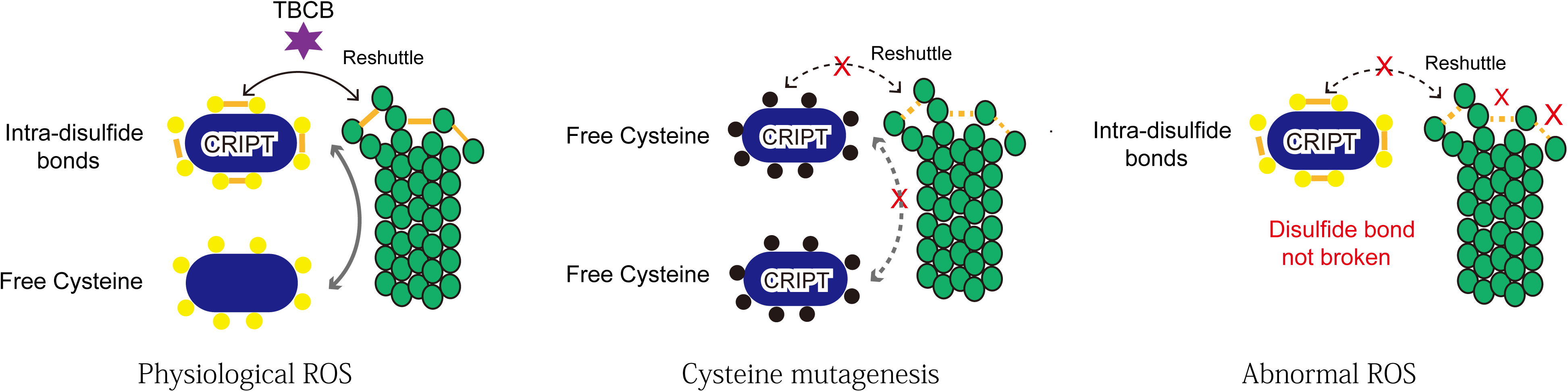
The model of CRIPT-controlled spindle disassembly in human cells. During normal mitosis, CRIPT exchange its disulfide bonds with tubulin subunits to facilitate the spindle disassembly. When CRIPT has cysteines mutated or cells face a disordered ROS environment, the thiol/disulfide exchange is blocked, resulting in impair of microtubules disassembly.

### CRIPT is a nuclear protein in interphase cells

CRIPT is a highly conserved cysteine-rich small protein. Mammalian CRIPT was originally identified in 1998 as an interacting partner of postsynaptic protein PSD95 (Niethammer et al., 1998), and its roles in neurons have been investigated by several research groups since then (Passafaro et al., 1999; Zhang et al., 2017). The published data demonstrated CRIPT as a cytoplasmic protein whatever it is native or exogenously expressed. However, it was found in our study to have an exclusive nuclear localization in neurons and other non-dividing cells. To reconcile these apparently conflicting observations, we hired different approaches and took a great care to determine its cellular localization. All our data from both biochemical and morphological assays clearly showed that CRIPT was restricted in nucleus with a remarked nucleolar concentration in interphase cells. Naturally, the announcements from the previous studies that CRIPT anchors synaptic receptors or promotes the growth of neuritis should be reconsidered. Although we cannot exclude the possibility that CRIPT could be expressed in the cytosol at the certain physiological or pathological conditions, it is more likely a nuclear rather than cytoplasmic protein inferred from its amino acid composition and the evolutionary origin. CRIPT is a very basic protein containing four typical CXXC motifs which are common in transcription factors and other DNA/RNA binding proteins. There is some data in public database supporting it as a nuclear protein in mammalian cells, including (1) Images of cell staining from the Human Protein Atlas using two different antibodies exhibit nuclear/nucleolar localizations of CRIPT in human and mouse cell lines (https://www.atlasantibodies.com/products/antibodies). (2) Protein similarity research reveals human CRIPT has two paralogs, and they are nucleolar proteins: RNA polymerase subunits POLR1C and POLR2C. (3) Most of the current known CRIPT interacting partners identified by affinity-captured MS are DNA or RNA binding nuclear proteins, such as THOC2, ELAVL1, zbtb48, MED23. Combining these public data with our experimental results, we here re-annotate CRIPT as a nuclear protein. The concentrated nucleolar localization is an agreement with its status as a paralog of RNA polymerase subunits, therefore, CRIPT would be functionally linked to RNA or ribosomal biogenesis in interphase cells. The nucleolus is best known as a housekeeping structure for ribosomal RNA production and ribosome assembly, however, accumulating evidence has revealed its roles much beyond ribosome biogenesis including cell cycle regulation (Visintin and Amon, 2000). Our present studies that have identified CRIPT as a key factor determining the fate of mitotic spindles have further highlighted the importance of nucleolus in cell cycle regulation.

### CRIPT senses and transfers intracellular redox signals to mitotic spindles

Recent studies have established that cell cycle is tightly coordinated with intracellular redox oscillations (Conour et al., 2004). The tubulin dimer has 20 cysteine residues, and can be oxidized by peroxynitrite *in vitro* and *in vivo* (Landino et al., 2007). Differently to other covalent modifications, redox reactions can be highly dynamic in physiological conditions and could mediate a switch-like response to fit the requirement of swift turnover of mitotic spindles during cell cycle. In this study, we for the first time observed the evidence that the disassembly of mitotic spindles is governed by a redox-dependent mechanism in which the cysteine-rich CRIPT protein transfers intracellular redox signals to spindles via thiol/disulfide exchange. This finding would be an important breakthrough in the field of cell cycle research.

As a novel mechanism identified for spindle disassembly, many issues related to redox regulation are needed to be addressed. Firstly, which and how redox signals are sensed and reacted by CRIPT-tubulin partners is waiting to answer. Cysteine thiols in proteins can be oxidized by proximal oxidases and intracellular enhanced ROS. Given the spindle movements are abrupt and robust changes during the cell cycle, we speculate that oxidation modifications on CRIPT-tubulin are most likely exerted by ROS rather than enzymatic mechanisms. ROS oxidation of cysteine thiols in cytoplasmic proteins is generally disfavored for both thermodynamic and kinetic reasons (Britto et al., 2005), therefore, cytoplasmic redox reactions are highly limited and selective. Although all of cysteine residues in tubulin subunits are structurally accessible to oxidative reagents, they seem to have no capability to directly sense and react with cytoplasmic ROS but are oxidized by CRIPT-bearing redox signals because redox-deficient CRIPT yielded highly stable mitotic spindles. The cysteine thiols in CRIPT were shown to have higher reactivity to H_2_O_2_ than those in tubulin in our *in vitro* assays (unpublished data), and this difference of thiol reactivity may account for the reason why CRIPT but not tubulin senses the intracellular ROS. The intracellular oxidants are known to preferentially react with deprotonated nucleophilic thiolates, and most cysteine thiols are protonated at physiological pH, rendering them not susceptible to oxidative reaction (Rudyk and Eaton, 2014). In addition to the pH, thiol reactivity is also determined by the electrostatics of the local environment andthe accessibility of the cysteines. Although CRIPT has a very high overall pI, the pKa of thiols in first four cysteines could be lowered by proximity to proton accepting amino acids and would be susceptible to oxidation, and the thiol in Cys3 would be more preferentially oxidized due to its extremely terminal localization. This speculation is supported by our result that the first two CXXC pairs are major targets in CRIPT redox reactions.

Secondly, how redox signals are transferred from CRIPT to tubulin subunits is needed to be resolved. Cysteine oxidation of tubulin is thought to be accompanied by loss of polymerization competence (Britto et al., 2005). However, the modification is unlikely to occur only on few of cysteines to achieve MT depolymerization *in vivo* because the formation of one or two disulfides was actually shown to promote polymerization (Chaudhuri et al., 2001). Our result that each of cysteine substitution in CRIPT all caused the defects of spindle movements highly suggests that the thiol/disulfide exchange with tubulin should occur at a multiple level. Differently to a single thiol/disulfide exchange which is relatively straightforward, the avenues of the multiple thiol/disulfide exchanges must be much more complicated. Analogous to multisite phosphorylation on a protein which can be distributive or progressive, the multiple thiol/disulfide exchanges between two molecules may occur orderly or stochastically up to a threshold number to accomplish a collective effect. In the case of CRIPT-tubulin, the redox exchanges could undergo at an ordered site-to-site sequence. Human tubulin subunits have acidic pI values ranging from 5.2 to 5.8, thus, CRIPT-tubulin associations could be electrostatic interaction. However, our result showed that all of the information for their interaction could be present in the primary amino acid sequence because their direct interactions were largely dependent on both GKNKF sequence and intactness of cysteine residues in CRIPT. Therefore, the direct contacts and redox exchanges between CRIPT and tubulin subunits could be strictly orientated. Each of cysteine mutation in CRIPT yielded long-lasting spindles, seemingly meaning that all cysteines undertake redox signaling as a whole or at one-layered level. However, there were two types of long-lasting spindles (regular or multipole spindles) linking to the defects of CRIPT redox competence. Multipole spindles were only caused by the mutants with substitution in last four cysteines (CRIPT^L4^), indicating that the cysteines in CRIPT function at multi-layered level. *In vitro* biochemistry assays also revealed these cysteines have distinct sensitivity to H_2_O_2_ (unpublished data). Therefore, both functionality and redox reactivity of cysteines in CRIPT are hierarchical.

### CRIPT may serve as a target for cancer therapy

Our findings not only revealed a novel mechanism that mammalian cells employ to disassemble their spindles after telophase but also unmasked a fragile site in dividing cells where a defect in a key molecule would completely arrest the progression of cell division. Thus, CRIPT or related components in this site could serve as a substrate to develop the targeted drugs for cancer therapy. Nowadays, in spite of multiple modalities being applied in clinic, MT stabilizers such as taxol still remain a cornerstone in the treatment of many types of cancers (Jordan and Wilson, 2004). Functionally, cysteine-mutated CRIPT is a taxol-like MT stabilizer with a specificity restrictive to mitotic spindle, representing an ideal prototypic drug for cancer treatment. Attributing to its small molecular size and net positive charge, cysteine-mutated CRIPT proteins can be easily delivered into cells as a small cargo conjugated to a cell penetrating peptide (Betty et al., 2017) or directly enter cells as a potential cell penetrating protein. Meanwhile, as a gene product, therapeutic CRIPT can be specifically targeted and expressed in certain cancer cells. These advantages as therapeutic molecules have persuaded us to test their anti-tumor effects, and our preliminary data in mice models demonstrated that xenograft subcutaneous tumors underwent a rapid hemorrhage and necrosis when they were locally administrated with the exosomes containing vectors expressing CRIPT mutants (Figure S2). In addition to the utilization of CRIPT proteins, CRIPT related redox components could also be used as targets to develop small molecular compounds against the progression of cell division and cancer growth.

## Material and Methods

### Cell culture and antibodies

HeLa and Embryonic kidney HEK293 cell lines were cultured in DMEM medium supplemented with 10% FBS. Commercial primary antibodies used for western blotting, immunoprecipitation and immunofluorescence were mouse monoclonal antibody anti-α-tubulin (ab18251; Abcam), anti-β-tubulin (ab15568; Abcam), anti-γ-tubulin (ab179503; Abcam) and anti-β-actin (ab8227; Abcam), rabbit monoclonal antibody anti-biotin (ab234284; Abcam), rabbit polyclonal antibody anti-GST (ab19256; Abcam) and anti-His (ab9108; Abcam), respectively. Monoclonal anti-CRIPT antibody used for immunofluorescence of endogenous CRIPT was purchased from Altlas (HPA046080 and HPA056765). Anti-CRIPT rabbit polyclonal antibody was also generated in-house against full-length purified CRIPT protein. Goat anti–mouse secondary antibody IgG-HRP (ab97051, Abcam) and anti–rabbit IgG-HRP (ab205719, Abcam) were used in immunoblotting. Goat anti-mouse IgG H&L conjugated with Alexa Fluor 488 (ab150113, Abcam) and cy3 (ab97035, Abcam) were used as secondary antibodies for immunofluorescence.

### DNA constructs and knockout experiments

The *DsRed* gene sequence purchased from Sangon Biotech was cloned into pcDNA3.1 (Invitrogen) using NheI and BamHI to generate DsRed-fusioned vector pcDNA-DsRed. Human *cript* gene sequence was purchased from Sangon Biotech and cloned into pcDNA-DsRed using BamHI and HindIII to generate pcDNA-DsRed-*cript*. Human *cript* gene was also cloned into pEGFP-N1 (Clontech) using XhoI and BamHI to generate pEGFP-*cript*. The *cript* mutations and deletions were constructed using the Quick-Change site-directed mutagenesis kit (Stratagene) based on pcDNA-DsRed-*cript* according to the manufacturer’s protocol. The coding sequences of the potential CRIPT partner were amplified using PCR from human cDNA. The fragments were then inserted into pGEX-6p-1 expression vector to generate GST-tagged proteins using BamHI and XhoI for β-actin, TBCB, ACTR2, ARPC1B and CAPZB; EcoRI and XhoI for α-tubulin, β-tubulin and TPM3, respectively. All restriction enzymes were purchased from NEB (New England Biolabs). The *cript* and *tbcb* gene knock out experiment was performed on HeLa cell lines using CRISPER-Cas9 system with gDNA sequence: KOct1: 5’-CACCGCACCATCTTTCCATGTATC-3’, KOct2: 5’-AAACGATACATGGAAAGATGGTG-3’ and KOtb1: 5’-CACCGTATCGCTTCTCGGAGCGGA-3’, KOtb2: 5’-AAACTCCGCTCCGAGAAGCGATACGGTG-3’, respectively (https://zlab.bio/guide-design-resources). Plasmid transfections were performed using the calcium phosphate method in both HeLa and HEK293T cells.

### Immunofluorescence and live-cell imaging

HEK 293 or HeLa cells grown on poly-lysine coated coverslips were washed with PBS for three times and fixed with 4% paraformaldehyde at 4°C for 15min, followed by permeabilized with 0.5% Tween X-100 in PBS for 20 min. After rehydration with PBS for 10 min and blocking for 1 h in 5% goat serum in PBS at room temperature (RT), coverslips were incubated with primary antibodies diluted in blocking buffer at 4°C overnight. Cells were then incubated with secondary antibodies conjugated with Alexa Fluor 488 or cy3 for 1 h at RT. After three washes with 0.05% Tween-20 in PBS, DNA was counterstained with DAPI (D1306; Invitrogen). Coverslips were mounted with FluorSave reagent (345789; Merck). Images were obtained using a confocal fluorescence microscope FV1200MPE-share (Olympus) equipped with a 60× Plan Achromat oil immersion objective (NA 1.25) at RT. Images were acquired using FLU OVI EW viewer software (Ver. 4.2; Olympus). For time-lapse microscopy, HeLa cells expressing CRIPT^WT^ or CRIPT^C3Y^ were cultured on glass-bottom 35 mm culture dishes (NEST Instruments) and assembled in a stage-top incubator supplying with 5% CO_2_. Sequences of images were acquired every 15 min for 24 hours using an inverted fluorescence microscope (Olympus) equipped with a 25× Plan Achromat oil immersion objective (NA 1.25) and a cooled charged coupled device camera (QImaging). Images were analyzed using Image-Pro Plus.

### Western blotting, immunoprecipitation, and GST pull down

After transfected for 48 hours, cells were harvested from 9 cm dishes by centrifugation at 2000 rpm for 5 min and then washed once with ice-cold PBS. The cells were lysed with lysis buffer (150 mM NaCl, 20 mM Tris-HCl, pH 8.0, 1% Triton X-100, 2 mM PMSF, 2 μg/ml leupeptin, 2 μg/ml aprotinin, 1 μg/ml pepstatin A, 1 mM Na_3_VO_4_, 5 mM NaF, and 10 mM β-glycerophosphate) and subjected to western blotting. For co-immunoprecipitation, the cell lysates were centrifuged at 12,000 g for 15 min at 4 °C and the supernatant was precleared with Pierce Protein A/G Magnetic Beads (88802, Thermo scientific) for 15 min on ice and then incubated with 2 μg anti-CRIPT antibody at 4 °C for 2 hours and with Pierce Protein A/G Magnetic Beads for another 2 hours. The beads were washed three times with lysis buffer and three times with ice-cold PBS before loading on SDS-PAGE. The gel was analyzed using silver-staining and the suspected band was excised for mass spectrometric analysis (Sangon Biotech). Expression of GST tagged β-actin, TPM3, ACRT2, CAPZB, TBCB and APRC1B were induced with 1mM IPTG in *E. coli* (BL21) at 37 °C for 4 h. Cells were lysed by sonication followed by centrifugation at 12,000 g for 15min. The supernatant was mixed with 50% slurry of glutathione sepharose 4B (GE healthcare) with gentle agitation at RT for 1 h. After centrifugation at 500 g, the supernatant was removed and the matrix was washed with PBS for three times. The matrix (GST-protein-beads) was finally mixed with purified His-tagged CRIPT with gentle agitation at 4 °C for 1 h. The beads were washed with PBS for three times before analysis by western blotting.

### Cysteine labeling assay

Purified tubulin or/and CRIPT at 1-2 mg/ml were dissolved in 0.1 M NH_4_HCO_3_ and incubated with H_2_O_2_ for 30 min at 22 °C. To remove excess H_2_O_2_, 500 U catalase was added in a total reaction volume of 20 µl. IAF (dissolved in DMF) was added to a 10-fold molar excess over protein cysteines and incubated for 2 hours in the dark at 22°C. The reaction was stopped by direct addition of loading buffer containing β-mercaptoethanol, and the labeled proteins were examined on SDS-PAGE and imaged by ECL system.

### Biotin switch assay

The biotin switch-labeling method was performed as follows. Briefly, transfected Hela cells were lysed in TNE buffer (1% NP-40, 50 mM Tris-HCl, pH 8.0, 140 mM NaCl, 5.0 mM EDTA and 1.0 mM NaF) in the presence of 1mM Na_3_VO_4_, 1mM PMSF and protease inhibitor mixture. Free thiols in proteins were blocked with 15 mM of N-ethylmaleimide (NEM) in the presence of 2.5% (w/v) SDS and incubated for 20 min at 50°C. Unreacted NEM was removed using a desalt spin column, and protein samples were eluted in TNE buffer containing 0.5% SDS. Disulfide bonds were converted to free thiols with 10 mM DTT. The free thiols were then labeled with the biotin-linked reagents N-biotinylaminoethyl methanethiosulfonate (MTSEA-biotin) (0.15 mg/ml) at 25 °C for 1 h. MTSEA-Biotin only reacts with cysteinyl sulfhydryl groups and was used to label free cysteines in protein. Biotinylated proteins were captured with streptavidin-sepharose (GE Healthcare) and then analyzed by western blotting.

## Acknowledgement

This work was supported by the National Natural Science Foundation of China (81572364 to C. Cai) and the Natural Science Foundation of Anhui Province (1608085MH222 to C. Cai).

## Author Contributions

Kehan Xu, Hong Li and Chunlin Cai conceived the project. All authors performed the experimental work. Kehan Xu, Chunlin Cai analyzed the data and prepared figures. Kehan Xu, Hong Li and Chunlin Cai wrote and edited the manuscript. Chunlin Cai supervised all aspects of the project.

---

**Figure S1.**
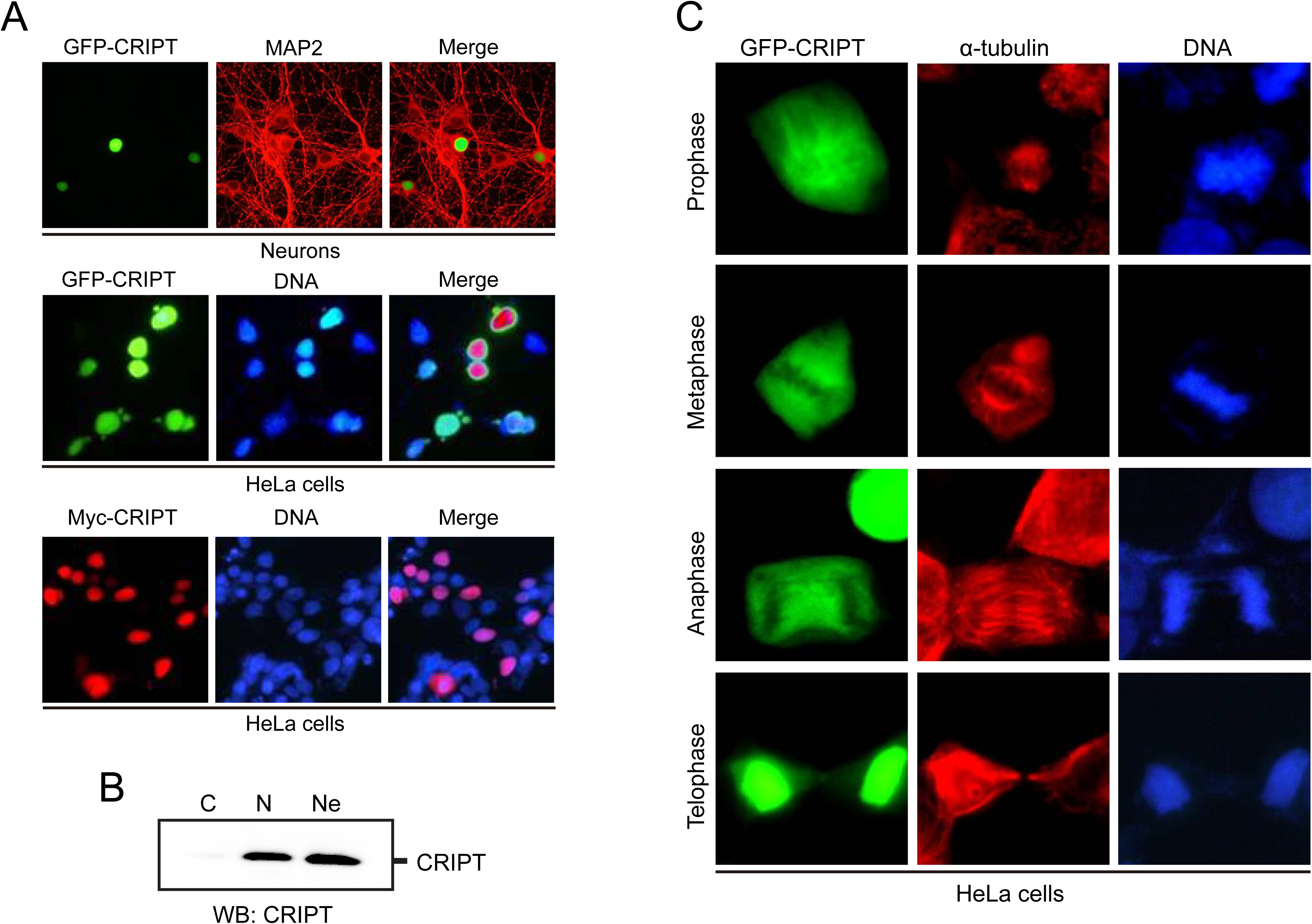
CRIPT is a nuclear protein and colocalizes with α-tubulin during mitosis. **(A)** Top and middle panel: GFP tagged CRIPT protein was expressed in both neurons and HeLa cells. MAP2 protein was immune-stained to visualize neurons (top panel). Bottom panel: HeLa cells expressed Myc-CRIPT was immune-stained with anti-myc monoclonal antibody. DNA was counterstained with DAPI. Scale bar, 10 μm. **(B)** Subcellular distribution of CRIPT protein was examined by cell fractionation. C: cytoplasmic, N: nucleus, Ne: nucleolus. **(C)** Co-localization between GFP-CRIPT and α-tubulin throughout mitosis.

**Figure S2.**
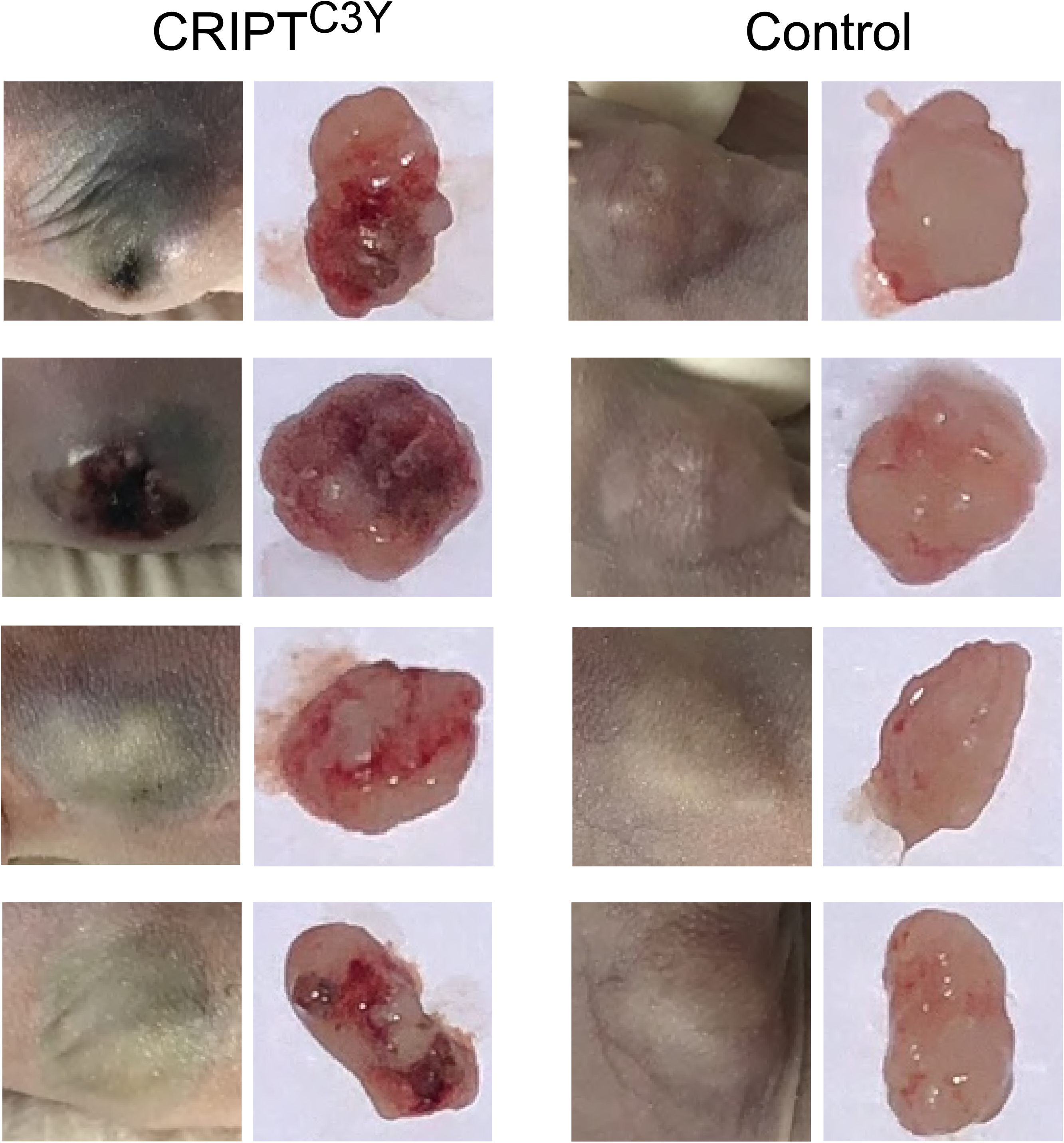
The anti-tumor effect of the mutated CRIPT protein. The mice subcutaneous tumors grown from MGC ells were locally injected with the exosomes containing vectors expressing CRIPT mutants or parental vector as control. The pictures were taken at 5 days after injection. Left panel shows the tumors treated with vectors expressing CRIPT^C3Y^, while right panel shows the tumors treated with control vector.

**Video 1**. HeLa cells expressing DsRed-CRIPT^WT^ display the normal cell division.

**Video 2**. HeLa cells expressing DsRed-CRIPT^C3S^ show the formation of Type A spindles with two bright spots at poles.

**Video 3**. HeLa cells expressing DsRed-CRIPT^C3S^ show the formation Type B messy fiber-like spindles.

## References

Asbury, C.L. (2017). Anaphase A: Disassembling Microtubules Move Chromosomes toward Spindle Poles. Biology (Basel) 6, 15.

Betty, R.L., Yue-Wern, H., Mallikarjuna, K., Shih-Yen, L., Robert, S.A., and Han-Jung, L. (2017). The Primary Mechanism of Cellular Internalization for a Short Cell-Penetrating Peptide as a Nano-Scale Delivery System. Current Pharmaceutical Biotechnology 18, 569–584.

Britto, P.J., Knipling, L., McPhie, P., and Wolff, J. (2005). Thiol-disulphide interchange in tubulin: kinetics and the effect on polymerization. Biochem J 389, 549–558.

Brown, M.R., Miller, F.J., Li, W.-G., Ellingson, A.N., Mozena, J.D., Chatterjee, P., Engelhardt, J.F., Zwacka, R.M., Oberley, L.W., Fang, X., et al. (1999). Overexpression of Human Catalase Inhibits Proliferation and Promotes Apoptosis in Vascular Smooth Muscle Cells. Circulation Research 85, 524.

Chaudhuri, A.R., Khan, I.A., and Ludueña, R.F. (2001). Detection of Disulfide Bonds in Bovine Brain Tubulin and Their Role in Protein Folding and Microtubule Assembly in Vitro: A Novel Disulfide Detection Approach. Biochemistry 40, 8834–8841.

Conour, J.E., Graham, W.V., and Gaskins, H.R. (2004). A combined in vitro/bioinformatic investigation of redox regulatory mechanisms governing cell cycle progression. Physiological Genomics 18, 196–205.

Ibarlucea-Benitez, I., Ferro, L.S., Drubin, D.G., and Barnes, G. (2018). Kinesins relocalize the chromosomal passenger complex to the midzone for spindle disassembly. The Journal of cell biology 217, 1687–1700.

Jones, D.P. (2010). Redox sensing: Orthogonal control in cell cycle and apoptosis signaling. Journal of internal medicine 268, 432–448.

Jordan, M.A., and Wilson, L. (2004). Microtubules as a target for anticancer drugs. Nature Reviews Cancer 4, 253–265.

Landino, L.M., Koumas, M.T., Mason, C.E., and Alston, J.A. (2007). Modification of Tubulin Cysteines by Nitric Oxide and Nitroxyl Donors Alters Tubulin Polymerization Activity. Chemical Research in Toxicology 20, 1693–1700.

Landino, L.M., Moynihan, K.L., Todd, J.V., and Kennett, K.L. (2004a). Modulation of the redox state of tubulin by the glutathione/glutaredoxin reductase system. Biochemical and biophysical research communications 314, 555–560.

Landino, L.M., Robinson, S.H., Skreslet, T.E., and Cabral, D.M. (2004b). Redox modulation of tau and microtubule-associated protein-2 by the glutathione/glutaredoxin reductase system. Biochemical and biophysical research communications 323, 112–117.

Laurent, A., Nicco, C., Chéreau, C., Goulvestre, C., Alexandre, J., Alves, A., Lévy, E., Goldwasser, F., Panis, Y., Soubrane, O., et al. (2005). Controlling Tumor Growth by Modulating Endogenous Production of Reactive Oxygen Species. Cancer Research 65, 948.

Leduc, M.S., Niu, Z., Bi, W., Zhu, W., Miloslavskaya, I., Chiang, T., Streff, H., Seavitt, J.R., Murray, S., Eng, C., et al. (2016). CRIPT exonic deletion and a novel missense mutation in a female with short stature, dysmorphic features, microcephaly and pigmentary abnormalities. American journal of medical genetics Part A 170, 2206–2211.

Lee, S.E., Frenz, L.M., Wells, N.J., Johnson, A.L., and Johnston, L.H. (2001). Order of function of the budding-yeast mitotic exit-network proteins Tem1, Cdc15, Mob1, Dbf2, and Cdc5. Current Biology 11, 784–788.

Lewis, S., Wang, D., and Cowan, N. (1988). Microtubule-associated protein MAP2 shares a microtubule binding motif with tau protein. Science 242, 936–939.

Ludueña, R.F., Roach, M.C., Jordan, M.A., and Murphy, D.B. (1985). Different reactivities of brain and erythrocyte tubulins toward a sulfhydryl group-directed reagent that inhibits microtubule assembly. Journal of Biological Chemistry 260, 1257–1264.

Mellon, M.G., and Rebhun, L.I. (1976). Sulfhydryls and the in vitro polymerization of tubulin. The Journal of cell biology 70, 226–238.

Menon, S.G., and Goswami, P.C. (2006). A redox cycle within the cell cycle: ring in the old with the new. Oncogene 26, 1101.

Menon, S.G., Sarsour, E.H., Spitz, D.R., Higashikubo, R., Sturm, M., Zhang, H., and Goswami, P.C. (2003). Redox Regulation of the G_1_ to S Phase Transition in the Mouse Embryo Fibroblast Cell Cycle. Cancer Research 63, 2109.

Mishima, M., Maesaki, R., Kasa, M., Watanabe, T., Fukata, M., Kaibuchi, K., and Hakoshima, T. (2007). Structural basis for tubulin recognition by cytoplasmic linker protein 170 and its autoinhibition. Proceedings of the National Academy of Sciences of the United States of America 104, 10346–10351.

Niethammer, M., Valtschanoff, J.G., Kapoor, T.M., Allison, D.W., Weinberg, R.J., Craig, A.M., and Sheng, M. (1998). CRIPT, a Novel Postsynaptic Protein that Binds to the Third PDZ Domain of PSD-95/SAP90. Neuron 20, 693–707.

Passafaro, M., Sala, C., Niethammer, M., and Sheng, M. (1999). Microtubule binding by CRIPT and its potential role in the synaptic clustering of PSD-95. Nature Neuroscience 2, 1063.

Rhee, S.G. (1999). Redox signaling: hydrogen peroxide as intracellular messenger. Experimental & Molecular Medicine 31, 53.

Rudyk, O., and Eaton, P. (2014). Biochemical methods for monitoring protein thiol redox states in biological systems. Redox Biol 2, 803–813.

Sarsour, E.H., Agarwal, M., Pandita, T.K., Oberley, L.W., and Goswami, P.C. (2005). Manganese Superoxide Dismutase Protects the Proliferative Capacity of Confluent Normal Human Fibroblasts. Journal of Biological Chemistry 280, 18033–18041.

Schieber, M., and Chandel, N.S. (2014). ROS function in redox signaling and oxidative stress. Current biology : CB 24, R453–R462.

Shaheen, R., Faqeih, E., Ansari, S., Abdel-Salam, G., Al-Hassnan, Z.N., Al-Shidi, T., Alomar, R., Sogaty, S., and Alkuraya, F.S. (2014). Genomic analysis of primordial dwarfism reveals novel disease genes. Genome Research 24, 291–299.

Tu, B.P., Kudlicki, A., Rowicka, M., and McKnight, S.L. (2005). Logic of the Yeast Metabolic Cycle: Temporal Compartmentalization of Cellular Processes. Science 310, 1152.

Visintin, R., and Amon, A. (2000). The nucleolus: the magician’s hat for cell cycle tricks. Current opinion in cell biology 12, 372–377.

Wang, H.P., Schafer, F.Q., Goswami, P.C., Oberley, L.W., and Buettner, G.R. (2003). Phospholipid Hydroperoxide Glutathione Peroxidase Induces a Delay in G(1) of the Cell Cycle. Free radical research 37, 621–630.

Wilson, C., and González-Billault, C. (2015). Regulation of cytoskeletal dynamics by redox signaling and oxidative stress: implications for neuronal development and trafficking. Frontiers in cellular neuroscience 9, 381–381.

Woodruff, J.B., Drubin, D.G., and Barnes, G. (2010). Mitotic spindle disassembly occurs via distinct subprocesses driven by the anaphase-promoting complex, Aurora B kinase, and kinesin-8. The Journal of cell biology 191, 795–808.

Woodruff, J.B., Drubin, D.G., and Barnes, G. (2012). Spindle assembly requires complete disassembly of spindle remnants from the previous cell cycle. Mol Biol Cell 23, 258–267.

Zhang, L., Jablonski, A.M., Mojsilovic-Petrovic, J., Ding, H., Seeholzer, S., Newton, I.P., Nathke, I., Neve, R., Zhai, J., Shang, Y., et al. (2017). SAP97 Binding Partner CRIPT Promotes Dendrite Growth In Vitro and In Vivo. eNeuro 4, ENEURO.0175-0117.2017.

Zimniak, T., Stengl, K., Mechtler, K., and Westermann, S. (2009). Phosphoregulation of the budding yeast EB1 homologue Bim1p by Aurora/Ipl1p. The Journal of cell biology 186, 379–391.

